# Single shot low-dose radiation durably and focally increases cortical excitability: a potential therapy for neuronal circuit disorders?

**DOI:** 10.64898/2026.08.04.742920

**Authors:** Wei Fan, Jannik Maier, Ting Fu, Fabian Langenbahn, Fabian Peter, Saleh Altahini, Dirk Cleppien, Stephanie Hehlgans, Julian Anthes, M. Bret Schneider, Hemmings Wu, John R. Adler, Michael J. Schmeisser, Franz Rödel, Albrecht Stroh

**Affiliations:** Leibniz Institute for Resilience Research; Mainz, Germany; Institute of Neuropathology, University Hospital Münster; Münster, Germany; Institute of Anatomy, University Medical Center of the Johannes Gutenberg-University; Mainz, Germany; Institute of Physiology 1, University Hospital Münster; Münster, Germany; Department of Neurosurgery, Stanford University School of Medicine; Stanford, CA, USA; Zap Surgical Systems, Inc; San Carlos, CA, USA; Goethe University Frankfurt, University Hospital, Department of Radiotherapy and Oncology, Frankfurt am Main, Germany; Georg-Speyer-Haus, Institute for Tumor Biology and Experimental Therapy, Frankfurt am Main, Germany; Eberhards Karls University Tübingen, Department of Neurology and Interdisciplinary Neurooncology, Hertie Institute for Clinical Brain Research, Tübingen, Germany; Eberhard Karls University Tübingen, Cluster of Excellence iFIT (EXC2180) ‘Image guided and Functionally Instructed Tumor Therapies’, Tübingen, Germany

## Abstract

Herein, we assess the potential of low-dose stereotactic radiosurgery (SRS) to modulate neuronal network states without apparent damage to cellular integrity. Using a small animal radiation research platform (SARRP), a 1 mm^3^ focal target in the mouse visual cortex was irradiated with doses of 5, 20, and 40 Gy. One-month later a significant dose-dependent increase in excitatory synapse numbers was observed, notably limited to the treated visual cortex and not the adjacent somatosensory cortex. Six months post-irradiation, cortical neuronal microcircuit activity was monitored in awake mice using high sensitivity two-photon calcium imaging. A single 5 Gy dose resulted in a significant microcircuit-wide increase of spontaneous neuronal activity, consistent with a lasting shift in the functional architecture of the irradiated nodal network. At higher SRS doses (40 Gy) this neuromodulatory window appears to close. In aggregate, these data suggest that low-dose radiation could, in some circumstances, be exploited by selected high precision SRS technologies to durably modulate neuronal circuit disorders. Some, or even all the clinical benefits reported in the companion article by Zhao et al. are likely attributable to the biological properties we sought to characterize in our research.

**One Sentence Summary:** Low-dose stereotactic radiosurgery effectively and durably modulates neuronal excitability via synaptic re-organization and could open new clinical possibilities for neuromodulation.

## Introduction

Stereotactic radiosurgery (SRS) is an ionizing radiation procedure defined by: 1) the high spatial accuracy of targeting, and 2) the use of extreme beam crossfire which both together enable an exceptionally steep dose gradient thereby affording relative protection of normal surrounding anatomy. Over the past four decades radiosurgical techniques have evolved into a neurosurgical mainstay used primarily for “ablating” a gamut of small to medium sized brain lesions, the vast majority of which are benign or malignant neoplasms as well as various types of vascular malformations.

Based on the cumulative experience of having treated millions of patients, a broad spectrum of radiosurgical doses has been adopted depending on lesion histology, volume and anatomic location. Importantly, dosing has largely been driven by the seeming overarching “self-evident” therapeutic objective of target (mostly tumor) ablation, and in this regard, single fraction SRS doses have tended to range between 12 and 160 Gy ^1,2^. The most serious and common complication of brain SRS is unintended radionecrosis, the risk of which directly correlates in a complex manner with both radiation dose and treated volume. Nevertheless, it is widely accepted that the risk of radiation injury is minimal when the radiosurgical dose is less than 12 Gy and the target volume smaller than 5 cm^3^. As a point of reference, the radiosurgical “ablation” of a 1.6 cm diameter spherical brain tumor (volume of ∼2 cm3), might involve a marginal dose of 20 Gy (at lesional edge) as normalized to the 80% isodose line. In such a typical clinical scenario, the radiation dose when administered by a state-of-the-art photon radiosurgical device would, at a distance of 0.5 cm, fall below the 12 Gy threshold. Generally speaking, in a setting of high-quality SRS, small volume brain doses of less than 12 Gy have historically been deemed inconsequential.

Akin to radiofrequency and high intensity focused ultrasound surgery (HIFU) lesioning (e.g. anterior capsulotomies and VIM thalamotomies), higher dose radiosurgery (up to 160 Gy) can also be used, albeit less commonly, to make precise radionecrotic lesion for treating pain and functional neurological disorders, including amongst others, thalamotomy for chronic pain and essential tremor, as well as epilepsy^1,3^. Meanwhile, with doses ranging between 75-90 Gy, a relatively common application of SRS is the “treatment” of the retrogassarian trigeminal nerve in the setting of trigeminal neuralgia ^4^. Amongst such patients, a striking phenomenon has been occasionally observed whereby near immediate pain relief with preserved native sensory function (i.e. a neuromodulatory-like outcome) is experienced. Notwithstanding the relative frequency of trigeminal neuralgia radiosurgery, the underlying (radio)biology remains unexplained ^4^.

Despite a long history of high dose SRS being used as a non-invasive ablative surgical tool, there is mounting evidence for a new therapeutic window for SRS involving lower non-destructive dosing. Past studies in minipigs have revealed that “sub-ablative” doses of 40 Gy led to an increase in cortical excitability that persisted beyond six months post irradiation ^5^. These findings raise the possibility that sub-ablative SRS could have neuromodulatory value in a setting of neuronal circuit disorders such as depression and addiction. Herein, we asked, by using highly sensitive readouts of neuronal circuit function in a mouse model, whether the lower therapeutic window of non-ablative SRS might extend beneath 40 Gy. Recently, high-precision irradiation equipment for the treatment of very small target volumes in rodents has become commercially available. Such platforms include built-in computed tomography (image)-based positioning (IGRT: image guided radiotherapy), which when combined with a dedicated planning system, enable the delivery of spatially accurate “stereotactic” irradiation. The availability of such equipment permits for the first time the cellular effects of precisely focused radiation (i.e. radiosurgery) to be analyzed in clinically relevant (orthotopic) mouse models ^6^.

It is critical to point out that when seeking to neuromodulate a brain circuit and all its related functions, the preservation of core circuit functionality is of paramount importance. For low or ultra-low dose of SRS to meet this overarching objective, it means that any reset in network functionality must be both subtle and localized. The detection of such changes necessitates uniquely perceptive methods. Fortunately, neuronal activity at a cellular resolution is particularly amenable, at least in the mouse cortex, to highly sensitive optical observation methodologies, i.e. 2 Photon calcium imaging ^7–9^. Indeed, we and others have demonstrated, that *in vivo* 2-Photon calcium imaging allows for the early detection of sublet network state shifts in murine models of neurodegenerative disorders such as Alzheimer’s disease ^10,11^, Huntington’s disease ^12,13^, and Parkinson’s disease ^14^. In a mouse model of an episodic neuroimmunology disorder - multiple sclerosis -, a shift in network state persisting into remission was demonstrable, suggesting that maladaptive network states can outlive the initial pathophysiological event ^15,16^.

We hypothesize that 2 Photon calcium imaging could be a unique tool for studying the CNS effects of low-dose sub-ablative SRS. The overarching drive behind this research is the belief that with sufficient understanding single-shot SRS might, by virtue of producing a durable shift in the network state, represent a novel and potent non-invasive neuromodulation procedure with potential widespread clinical application among circuit disorders, such as addiction and treatment resistant depression. The conceptual framework for such a therapeutic approach is buttressed by recent advances in our understanding of the dynamics of network states which suggest an intrinsic stability of maladaptive, as well as adaptive states. This new model of network function seems to best explain how network attractors can be amenable to one-shot interventions, as for example, ketamine administration in the setting of TRD ^17^.

In the present set of experiments, we irradiated a small 1 mm^3^ target volume within the primary visual cortex of healthy mice with varying low dose SRS. One month after irradiation, confocal imaging and quantitative immunostaining was used to measure changes in well-characterized synaptic proteins that relate to excitatory and inhibitory synaptic transmission, both in the irradiated region, as well as in a neighboring cortical area. Six months later network function was assessed via 2 Photon calcium imaging. In summary these experiments demonstrate that beginning at only 5 Gy, there is a significant shift in network state without evidence of injury to the network integrity. Our results also suggest that synaptic re-organization is the most likely putative mechanism of action. The primary goal of this research was: 1) to investigate the threshold dose of radiation needed to induce the radiomodulation phenomenon, but even more importantly, 2) seek a better understanding of the basic neurophysiological mechanisms by which low dose irradiation alters both synaptic organization and local network function. In meeting these objectives, we intend to further elucidate the biological means that underlie the clinical results being reported in the Zhao et al companion article.

## Results

### High-precision irradiation of mouse visual cortex

Recent years have seen the emergence of innovative irradiation platforms that enable the accurate delivery of radiation to small discrete volumes within rodent brains. Utilizing such a small animal radiation research platform (SARRP), we irradiated specific regions of the mouse brain and subsequently studied the longitudinal effect on neuronal networks (Fig. 1A). Of note, built-in computerized tomographic (CT) imaging of individual mice allows specific anatomic landmarks to be accurately identified, which in conjunction with a mouse brain atlas, can establish the requisite stereotactic coordinates of the primary visual cortex prior to irradiation. (Fig.1 B to F). By subsequently using a focused two-beam radiosurgical-like geometry the visual cortex target could then be precisely irradiated somewhere between the 60% and 80% isodose lines. (Fig. 1E).

**Fig. 1.**
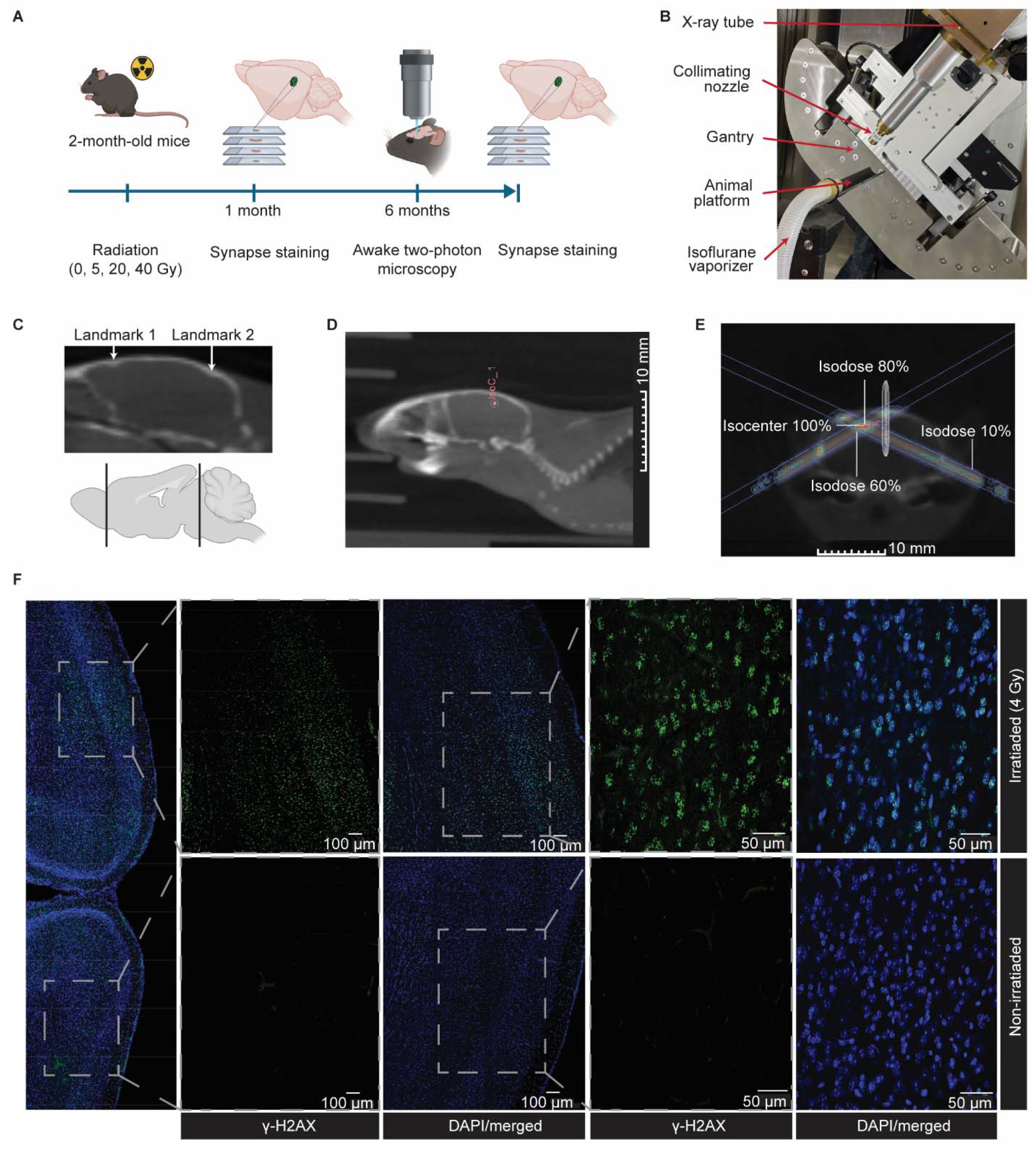
Stereotactic radiosurgery on mouse visual cortex. (**A**) Study design. (**B**) SARRP technical setup for irradiation treatment covering an X-ray tube mounted to a movable gantry allowing collimated dose delivery to mice located within an animal holder platform. (**C**) Treatment planning using anatomic landmarks accessible from built-in individual CT-scans. (**D**) Assessment of primary visual cortex coordinates in a mm scale based on the mouse brain atlas Isocenter (IsoC_1, 100% dose distribution) for precise dose targeting to the visual cortex. The isocenter was used to define the center to administer a two-beam irradiation geometry, using a 1 mm collimator. (**E**) Representative image of the dose distribution in the target region. The isocenter is depicted by the red dot. Color shades visualize the distribution of the relative dose. Shown is the frontal view including isodose 80%, 60% and 10 distribution (**F**) Images of γ-H2AX foci detection (green fluorescence) on cleared thick (300 μm) brain sections at 50 min after stereotactic targeted irradiation therapy with a dose of 4 Gy (upper panel). Control areas without irradiation are shown in the lower panel. DAPI was used for nuclear counterstaining. Scale bars indicate 100 μm and 50 μm.

To validate the technique used to irradiate such a small “millimetric” brain structure, *ex vivo* brain sections were stained for serine 139 phosphorylated histone H2A variant, γ−H2AX, a well- established marker of radiation damage and repair ^18^. Immunofluorescence detection of γ−H2AX confirmed the local enrichment of nuclear foci within the intended (planned) brain location (Fig. 1F, upper panel). By contrast, the absence or minimal detection of marker γ−H2AX in the non- irradiated control (lateral) hemisphere (Fig. 1F, lower panel) confirmed the targeting accuracy of our brain irradiation methodology.

### Irradiation leads to a dose-dependent shift in synapse densities towards higher excitability within the visual cortex

Initial experiments focused on cellular alterations that could potentially change the neuronal circuit function. Using immunostaining for synaptic markers that are associated with either excitatory or inhibitory synapses, we sought to quantify the impact of different single doses of radiation on excitatory-inhibitory synaptic organization (Figure 2). Shank2, a glutamate receptor-associated postsynaptic scaffold protein was used as a marker for excitatory synapses^19^, while γ- Aminobutyric acid type A receptor GABA_A_ (γ-2 subunit), a major postsynaptic inhibitory receptor of the CNS, was used to quantify the extent of inhibitory synaptic organization. In this set of experiments, both Shank2 and GABA_A_ puncta within the unilateral irradiated visual cortex were compared to the corresponding contralateral hemisphere one month after irradiation. Within the irradiated hemisphere a dose-dependent increase in Shank2 Puncta density was observed. (Figure 2).

**Fig. 2.**
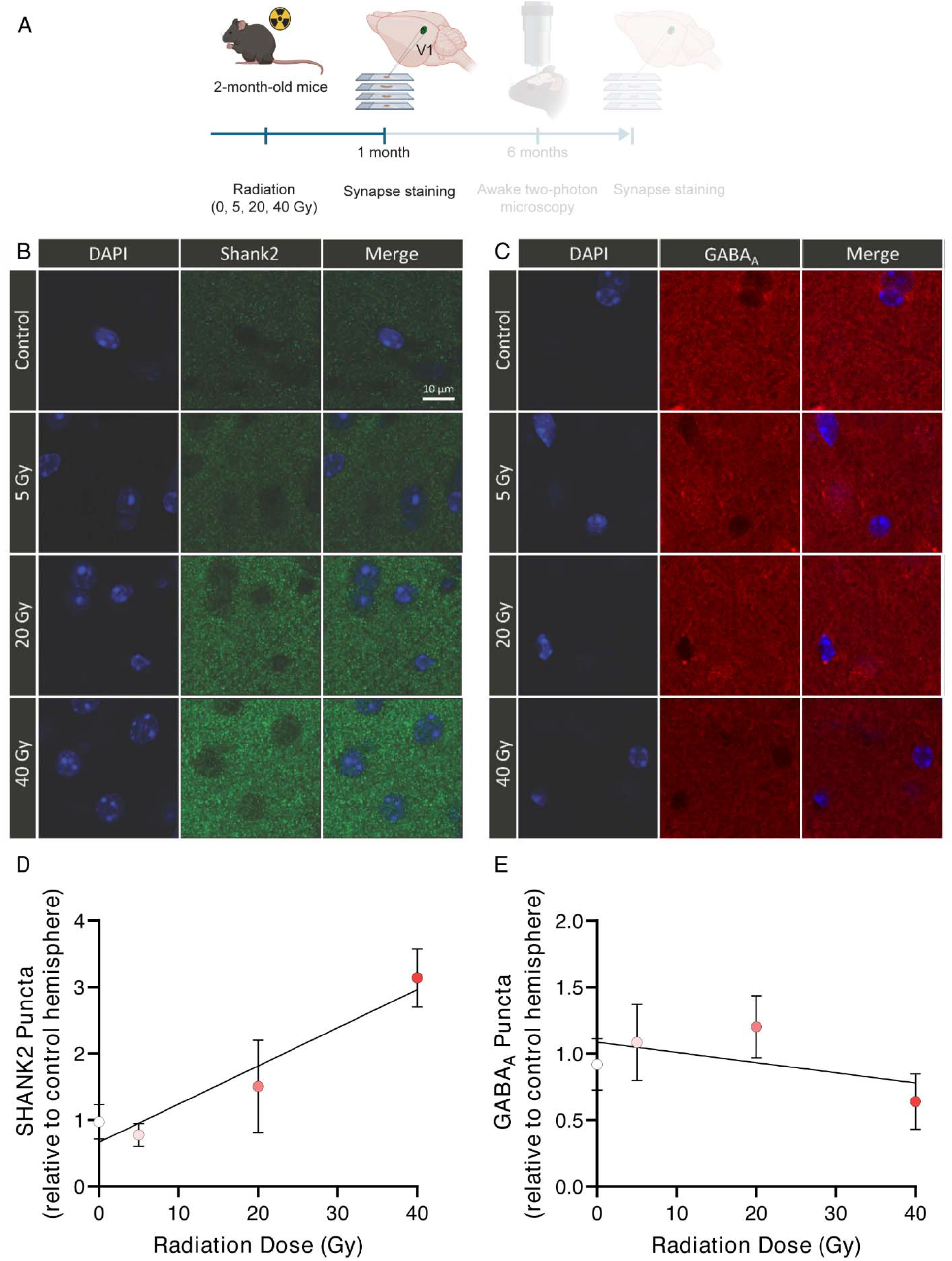
Radiomodulation alters postsynaptic scaffold protein organization within the visual cortex one month after treatment. **(A)** Experimental Design (B) Representative images of Shank2 puncta in control sections and sections one month after 5 Gy, 20 Gy, 40 Gy irradiation. **(B)** Representative images of GABAA receptor puncta in control sections and sections one month after 5 Gy, 20 Gy, 40 Gy irradiation. **(C)** Shank2 puncta quantification. Data were obtained from n = 5 animals per group with 8 replicates per animal. Correlation was analyzed by simple linear regression (Y = 0.05749X + 0.6622; slope = 0.05749; R² = 0.4885; F(1,18) = 17.19; p = 0.0006.) The slope is significantly different from zero. **(D)** GABAA puncta quantification. Data were obtained from n = 5 animals per group with 8 replicates per animal. Correlation was analyzed by simple linear regression (Y = −0.007677X + 1.086; slope = −0.007677; R² = 0.05428; F(1,18) = 1.033; p = 0.3229). The slope is not significantly different from zero. Data are presented as mean and s.e.m.

### Synaptic reorganization is restricted to the visual cortex, leaving the neighboring somatosensory cortex unaffected

To assess whether the Shank2-associated synaptic phenotype extended beyond the visual cortex into adjacent cortical regions, Shank2 puncta were quantified within the primary somatosensory cortex (Figure 3). In contrast to the primary visual cortex, quantification of Shank2 puncta revealed no statistically significant differences between irradiated and contralateral hemispheres at any irradiation dose (Figure 3C). Taken together, these findings indicate that the ionizing radiation-induced changes to Shank2 puncta density are anatomically restricted to a given irradiated brain circuit.

**Fig. 3.**
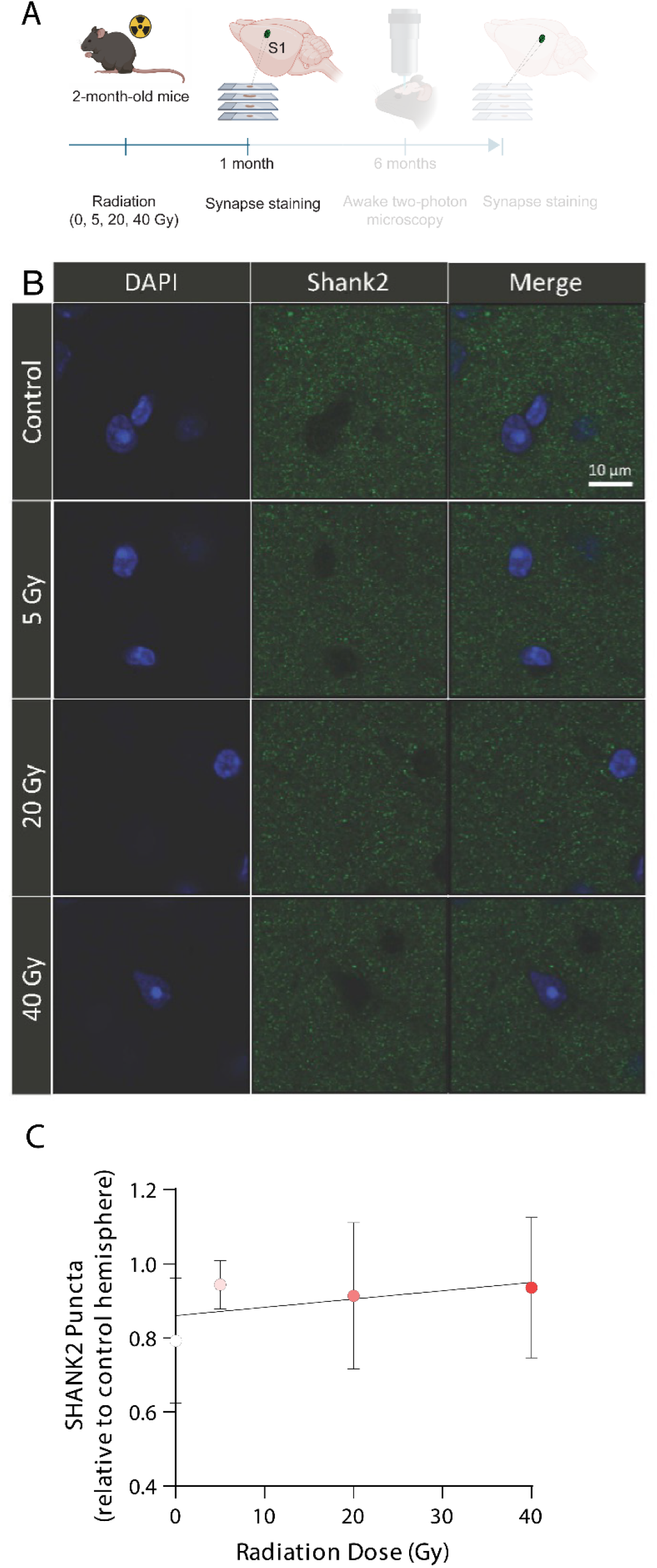
Irradiation induced altered Shank2 expression does not affect adjacent cortical regions, one month after irradiation. **(A)** Experimental Design (B) Representative images of Shank2 puncta in control sections and sections one month after 5 Gy, 20 Gy, 40 Gy irradiation in the somatosensory cortex. **(C)** Shank2 puncta quantification. Data were obtained from n = 5 animals per group with 8 replicates per animal. Correlation was analyzed by simple linear regression (Y = 0.002237X + 0.8601; slope = 0.002237; R² = 0.01086; F(1,18) = 0.1976; p = 0.662.) The slope is not significantly different from zero. Data are presented as mean and s.e.m.

### A single dose of 5 Gy significantly increased spontaneous neuronal firing at 6 months

Six months post SRS using doses of 5 Gy, 20 Gy and 40 Gy, 2-photon imaging was performed on awake mice. Non-irradiated animals served as controls. No change in cellular integrity was observed at any of the three dose levels. Regions of interest (ROIs) encompassing individual neuronal somata in layer II/III of mouse visual cortex at the area of irradiation were identified, and calcium transients were detected in a semi-automated manner (Fig. 4A). Binarized network activity of putative action potentials revealed a typical sparse firing in all four conditions, indicative of a healthy network ^20^. We did not observe increased network burstiness or hyperactivity (Fig. 4B). The portion of active cells were similar in all four groups (Fig. 4D, Fig. S1). Qualitative assessments of network synchronicity, as depicted by the correlation index matrix of paired ROIs, demonstrated that overall healthy modular synchronicity patterns were present under all four conditions (Fig. S2). These findings suggest, that all three doses of irradiation did not negatively impact the functional architecture of network function, a critical prerequisite for a clinical neuromodulation approach. A significantly increased (relative to the control mice) spontaneous firing rate was observed in the lowest 5 Gy cohort. (Fig. 2C). The measured increase of approximately 6%, is substantial given the otherwise tightly controlled cortical activity states. Increasing the SRS dose to 20 Gy further increased spontaneous firing rates by as much as 30%. However, at 40 Gy no significant increase in firing rate compared to control animals could be discerned. To summarize, the window of neuromodulation in our mouse model spanned from 5 to 40 Gy, with a peak at 20 Gy.

**Fig. 4.**
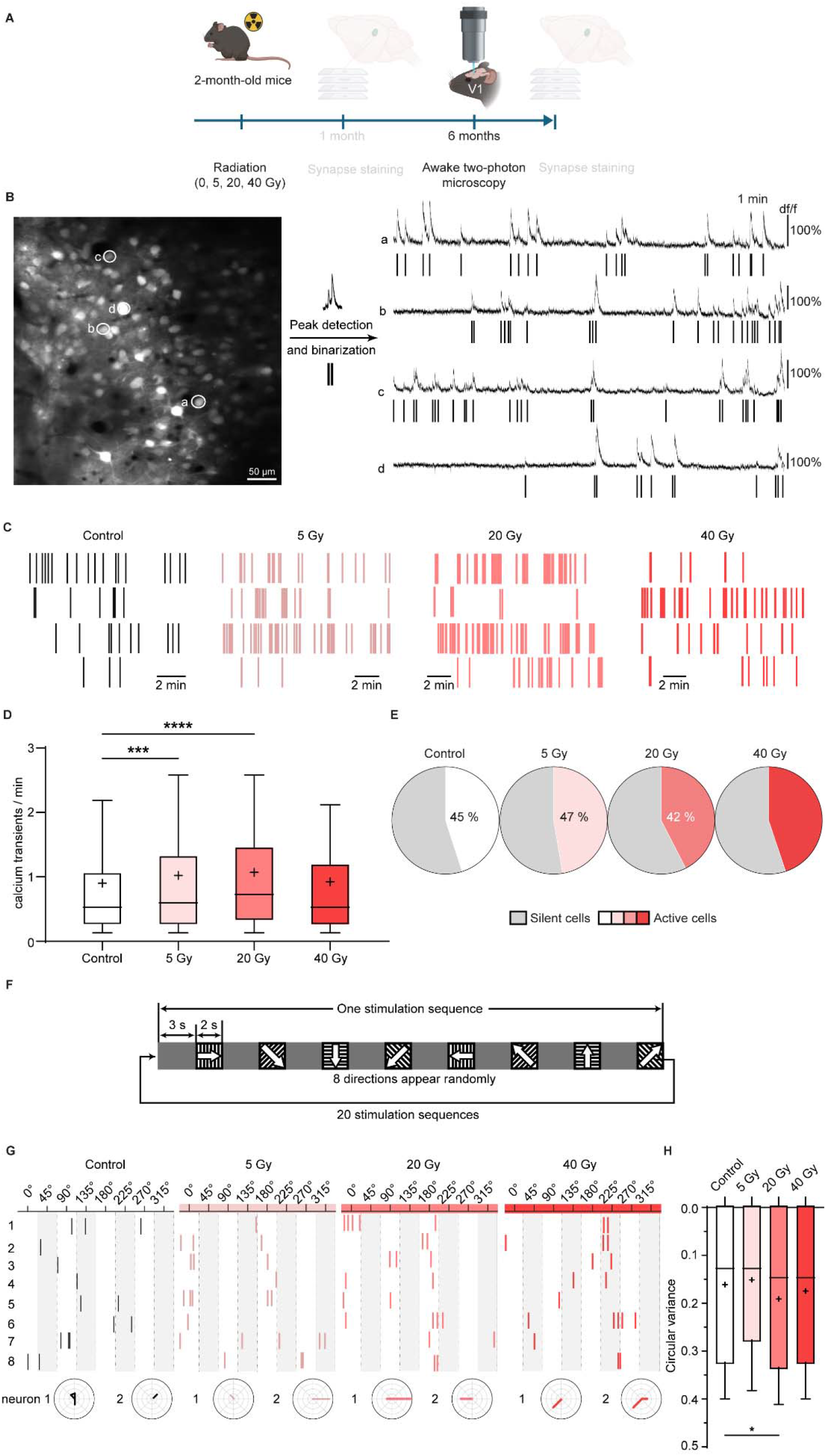
Single dose of 5 Gy irradiation increased spontaneous firing rate at 6 months after irradiation but did not change the direction selectivity of visually evoked responses to drifting gratings.. **(A)** Experimental Design **(B)** Workflow of calcium imaging analysis (**C**) Representative detected peaks after binarization. The average detected ROIs/mm^2^ ranged at more than 800. In total, 1350 ROIs from non-irradiated control group (7 mice), 3391 ROIs from 5 Gy irradiated group (10 mice), 2200 ROIs from 20 Gy irradiated group (8 mice) and 1240 ROIs from 40 Gy irradiated group (7 mice) were detected. (**D**) 5 Gy and 20 Gy irradiation increased spontaneous firing rate. (**E**) Irradiation did not affect the portion of active cells, as assessed by Kruskal-Wallis test. (**F**) Visual stimulation paradigm: in each stimulation cycle, after 3 seconds grey screen, 2 seconds drifting gratings appear in random order in 8 directions. One complete experiment comprised 20 stimulation sequences. (**G**) Representative binarized calcium transients (all responses of 20 stimulation sequences) and polar plots in response to visual stimulation from the control, 5 Gy, 20 Gy and 40 Gy irradiated mice. (**H**) Circular variance of 1114 ROIs from control group (7 mice), 2497 ROIs from 5 Gy irradiated group (10 mice), 1992 ROIs from 20 Gy irradiated group (8 mice) and 910 ROIs from 40 Gy irradiated group (6 mice).

### 5 Gy does not impact the directional selectivity to visual stimulation

To be appealing as a clinical procedure, a neuromodulation technique must preserve the primary functional integrity of any given cortical circuit. Hubel and Wiesel’s 1962 study demonstrated the elemental existence of orientation selective neurons in cat visual cortex ^21^. Seeking to assess the primary visual cortex (V1) function in SRS treated animals, each was subjected to the classical drifting grate paradigm that is based on Hubel and Wiesel’s pioneering study. Awake mice were presented with drifting gratings appearing randomly from eight directions (Fig. 4F) ^22,23^. We found that in all four conditions, neurons reliably responded to specific gratings, as indicated by highly selective response patterns in the polar plots (Fig. 4G). Selectivity of responses can be quantified by the circular variance as a global measure of the shape of the tuning curve, thereby characterizing the responses of sensory neurons to external stimuli ^24,25^. A dose of 5 Gy did not change the circular variance in comparison to the controls (Fig. 5H). However, a very slight decrease in visual tuning was observed at a radiation dose of 20 Gy, while firing rates among 40 Gy mouse cohort mice was similar to controls.

**Fig. 5.**
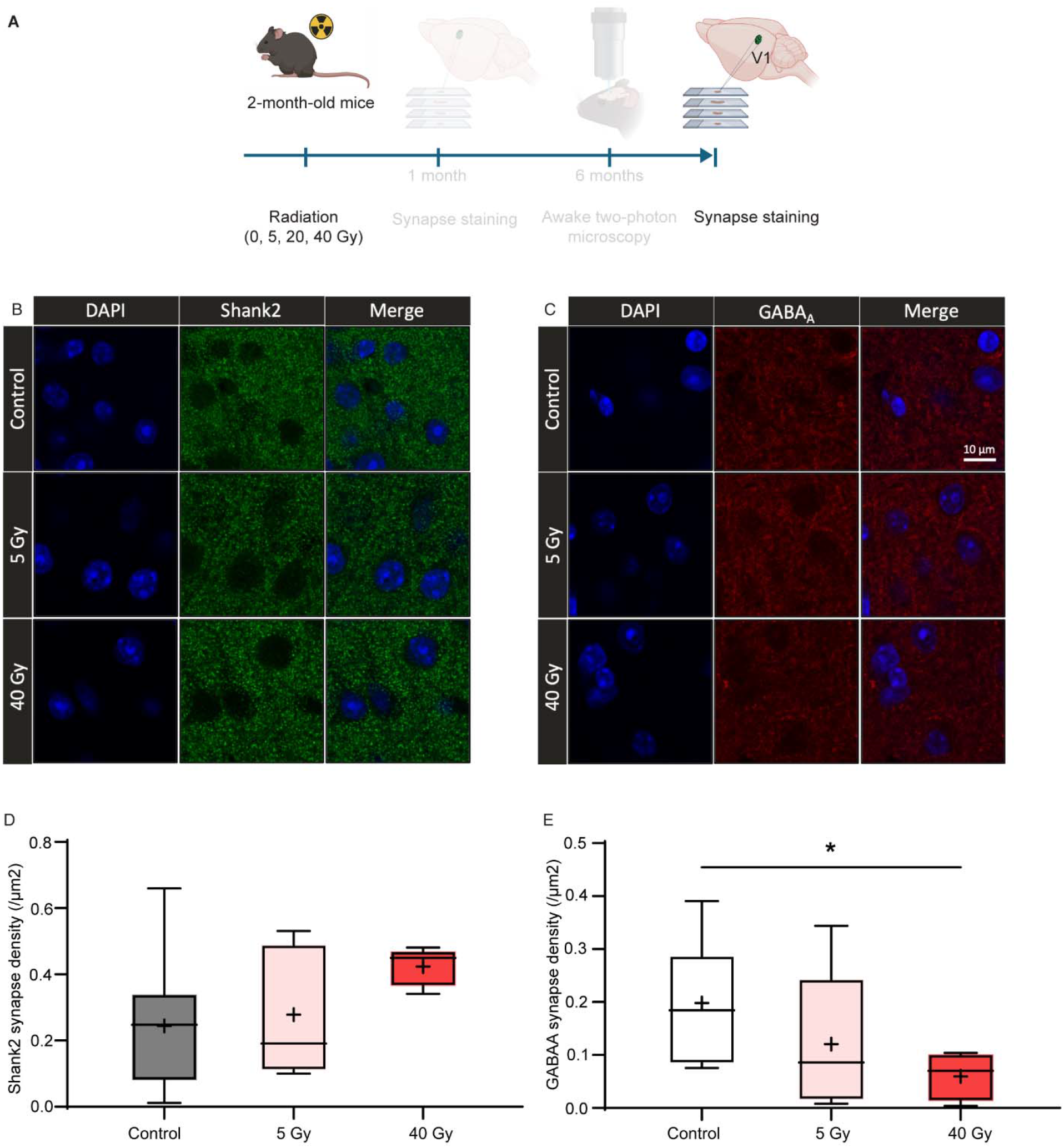
Irradiation affects excitatory-inhibitory synaptic organization within the visual cortex persisting after six months. (**A**) Experimental Design. (**B**) Representative Shank2 immunostaining and quantification count mask. (**C**) Representative GABAA immunostaining and quantification count mask. (**D**) 5 Gy or 40 Gy irradiation did not change the Shank2 synapse density (n = 10, 5 and 5 mice for the control, 5 Gy and 40 Gy groups). (**E**) 40 Gy irradiation decreased GABAA synapse density (n = 10, 5 and 5 mice for the control, 5 Gy and 40 Gy groups). Box and whiskers plot. Box indicates the 25- 75th percentile, whiskers indicate 10-90th percentile, and “+” indicates the mean. *p < 0.05. Kruskal-Wallis test.

### The shift in post-radiation excitation/inhibition persists for at least six months

The observed shift in excitatory and inhibitory synapse densities that was noted at one month post visual cortex irradiation was still observable at month six; such findings connect closely with parallel and near simultaneous assessments of functional microcircuit activity (Figure 5). While we observed a dose-dependent increase in the density of excitatory synapses, the density of inhibitory synapses showed a dose-dependent decrease (Fig: 5 E). The inhibitory synapse density and spontaneous firing rate changes observed in mouse visual cortex are consistent with an earlier study involving irradiated iPS-derived neuronal cultures, revealing an increased vulnerability of inhibitory neurons upon irradiaton ^5^.

## Discussion

### Could single shot low-dose focal radiation become a therapy for neuronal circuit disorders?

Our investigations have sought to elucidate the biological mechanisms through which low dose stereotactic radiation alters cortical neuronal dynamics. In a study involving focally irradiated healthy adult mice, highly sensitive functional readouts were combined with the measurement of markers of synaptic re-organization. Analogous to what was observed in the companion Zhao et al article, a powerful neuromodulatory effect could be demonstrated one month post SRS, albeit in the mouse experiments starting at a much lower dose; notably the 5 Gy dose threshold we found in mice was unknown to the designers of the TRD trial. Since this increase in spontaneous neuronal activity is still detected at 6 months, the effects of low dose radiomodulation, at least in mice, appear durable. Moreover, the preservation of precise representation of visual afferents after treatment with 5Gy makes localized radiation induced visual cortex damage improbable at this dose level; we are less certain that the same can be said for higher (20, 40 Gy) doses.

Our current findings in mouse visual cortex are clinically relevant in three ways: Firstly, by titrating a precise balance between excitatory and inhibitory synapses, and not inducing a radionecrotic brain lesion, low dose SRS opens up the intriguing prospect of being a potent new neuromodulatory technique for a range of presently incurable circuit disorders, including treatment resistant depression. Secondly, although the irradiation dose thresholds might not be identical in both rodent and human brains, a dose-dependent focal increase in the density of excitatory synapses in mouse visual cortex, and not in the neighboring somatosensory cortex, helps to address a possible safety concern prior to future human trials, demonstrating that the direct network alteration is restricted to the irradiated region. Lastly, the widespread but inadequately studied cognitive and neuropsychiatric symptoms that have been observed in patients undergoing large field brain irradiation (in the absence of brain injury on imaging) might now be reimagined to be caused by the neuromodulatory effects described herein.

How, in the absence of a brain lesion, might the effects of a single dose of ionizing radiation lead to a significant and durable shift in network excitability ^26^? This is particularly puzzling given that any sub-lethal effects of irradiation such as an increase in the oxidative stress level or protein and DNA damage by double-strand breaks should have been repaired within a few days at most after radiation exposure ^27–29^. While this topic merits further investigation, we hypothesize that synaptic plasticity, i.e. changing synaptic strength and modulating synaptic stability, as well as axonal re-organization, particularly in the grey matter of cerebral cortex, could all be key factors in the enduring effects of low dose SRS. For example, a somewhat analogous phenomenon may be involved in the longer-term benefits TDR patients experience undergoing a single ketamine administration ^30^. While the brain adapts to transient neuromodulatory stimuli over the short term, in doing so, the brain resets/learns over the longer term.

The trajectory of network function can be modelled and understood by dynamical systems theory ^17^. Stable network states can be conceptualized as attractors, i.e. states which are resistant towards smaller perturbations, ensuring network function in the face of adversity. The pursuit of stability, even in adaptive network states, e.g. a network state of depression, can be termed a “selfish network”, seeking to retain stability even if the phenotypic long-term outcome is detrimental to the organism^17^. Consequently, if single dose interventions such as low-dose SRS are strong enough to push the network trajectory to a different network attractor, in this case characterized by an increase in excitation, the new attractor is inherently stable, and can greatly outlast the duration of the treatment. The companion clinical article seemingly shows a similar newfound stability of the brain state setpoint in TRD patients.

### The merits of low-dose SRS relative to other current neuromodulation techniques

With the growing understanding of neuropsychiatric disorders being a consequence of dysregulated network dynamics, restorative therapies based on modulating key dysregulated hubs seem to be increasingly within reach^17^. Optogenetics has brought about a tremendous advance in our understanding of neuronal circuits in health and disease ^31^. However, optogenetics as a technique is currently, and quite possibly even inherently, limited to pre- clinical animal studies given the need to insert opsins into the neuronś genome, and then in a complicated second step, implant optical fibers for light delivery. Although neuromodulation with implanted deep brain stimulation (DBS) electrodes has been widely adopted to help manage symptoms in neurodegenerative disorders, e.g. Parkinsońs disease, its application to TRD and addiction are in its infancy, in large driven part by its intrinsically invasive nature.^32^. Transcranial magnetic stimulation (TMS) represents the most widely used non-invasive neuromodulation techniques for depression ^33,34^. Nevertheless, the physics underlying TMS make for rather ill-defined effects from the standpoint of spatial accuracy ^35^, particularly with respect to deep brain locations. Ultimately the benefit of TMS is modest and repeated interventions are the norm. Consequently, a single non-invasive intervention with permanent, or nearly so, clinical benefits, remains an elusive goal for the overall field of neuromodulation.

### Clear roadmap towards clinical implementation

As a biological and clinical phenomenon, we remain in the early stages of delineating the mechanisms underlying radiation-induced neuromodulation. Clearly more work is required to understand both the opportunities and the perils that such radiomodulation entails. However, the argument for treating human circuit-based disorders with low dose radiation has, by means of the companion article, moved beyond the realm of theory. In combining the findings from previous research with the current pair of articles, one is led to an inescapable conclusion; radiomodulation is a real biological phenomenon, the intrinsic properties of which, could enable an important set of neuromodulatory tools with broad potential clinical application.

In the accompanying first-in-human study involving patients with TRD, low-dose SRS was targeted at the subgenual anterior cingulate cortex, a key limbic structure that lay deep in the brain; the reported safety, rapidity of therapeutic onset and 3-month durability are all notable results of this treatment. Although the above is merely a pilot study, it is logical to speculate about other circuit-based disorders in which to radiomodulation’s effects might also be studied. For example, addiction could be a compelling next target: it is a circuit-based disorder of the same limbic and prefrontal networks, and notably, invasive DBS has shown growing efficacy in treating this disorder ^36^. Another unique aspect of radiomodulation is that high-dose precision radiation throughout the CNS has over the past generation become well established, albeit mostly in brain tumor patients, resulting today in hundreds of existing SRS facilities worldwide. Ultimately the convergent evidence presented in these companion articles warrants the significant expansion of careful sham controlled, randomized pilot, and even more comprehensive studies of radiomodulation across these and other neural circuit disorders.

## MATERIALS AND METHODS

### Study design

The aim of this study is to provide evidence for the prospect of using low radiation doses for long-term neuromodulation in a mouse model, as a reference for a future clinical study. To this end, we irradiated a small target volume in the visual cortex of 2-month-old mice with doses of 5, 20, and 40 Gy. Six months later, we monitored cortical neuronal microcircuit activity by 2- photon calcium imaging in the awake mouse. Spontaneous activities and visual stimulation responses are compared with the control groups. In the postmortem histology study, Shank2 and GABAA antibodies were used to quantify the excitatory and inhibitory synapse density, respectively.

### Animals

Female 2-month-old C57BL/6JRj were used in this study. All animals were group-housed at the animal facility of the Georg-Speyer-Haus, University of Frankfurt, for SARRP irradiation purpose and the Mouse Behavioral Unit Mainz, Johannes Gutenberg University Mainz and had ad-libitum access to food and water. All experimental procedures were performed in accordance with institutional animal welfare guidelines and were approved by the states of Hessen and Rhineland-Palatinate.

### Irradiation procedure

Irradiation was performed using the Small Animal Radiation Research Platform (SARRP, Xstrahl Ltd., Camberley, UK). Briefly, mice were anesthetized with isoflurane (1.5%) and imaged with an on-board cone beam computed tomography (CBCT) operating at 50 kV/0.8 mA and 720 projections acquired over 360°. Next, CBCT images were reconstructed using SARRP’s built-in Muriplan^TM^ planning software (Xstrahl Ltd) reconstruction algorithm into 0.16 mm isotropic voxels for contouring and individual isocenters were selected for targeted radiation therapy. Based on anatomical landmark 1 (the suture between the olfactory bulb and the cortex) and landmark 2 (lambdoid suture) accessible from CT scans, mouse brain stereotactic coordinates and the scale bars provided by the planning system, the primary visual cortex (coordinates from Bregma: left 2.5 mm, AP -2.7 mm (1.3 mm rostral from lambdoid suture), Z 0.5 mm) was targeted for precise irradiation. The isocenter (100% dose delivery) was used to define the center for administrating a two-beam irradiation geometry, using a 1 mm collimator and operating at 220 kV and 13 mA with a dose rate of 5.2 cGy/s. Mice received doses of 5 Gy, 20 Gy or 40 Gy, respectively, in a single fraction.

### Gamma-H2AX staining of brain slices

For immunofluorescence staining, paraformaldehyde (PFA) fixed brain samples were sliced into 300 μm thick sections using a Vibratome VT1200S (Leica, Nussloch, Germany). Brain slices were next cleared with a X-Clarity electrophoretic tissue clearing solution (Logos Biosystems, #C13001, Anyang-si, South Korea) at 0.6A and 37°C for 3 h. After tissue clearing, non-specific protein binding was blocked for 4 hours with 3% BSA in PBS containing 0.1% Triton-X100. Incubation of the primary antibody anti-phosphohistone H2AX (rabbit anti-γH2AX, Cell Signaling, Frankfurt, Germany, #9718, 1:200), an indirect marker of DNA double-strand break induction and repair ^18^ was performed in 1.5% BSA 0,1% TritonX-100 for 24 h at room temperature (RT) followed by incubation with fluorescent-labeled secondary antibodies (anti-rabbit AF488, Thermo Fisher Scientific, Darmstadt, Germany, #A11034, 1:500) overnight at RT. DAPI solution used at a 1:2500 dilution was applied for nuclear counterstaining (10 min, RT). Samples were embedded in X-Clarity mounting medium (Logos Systems #13101) and immunofluorescence staining on cleared thick sections was visualized applying a Yokogawa CQ1 confocal microscope (Yokogawa, Musashino, Japan), using either 20x or 40x objectives ^37^.

### Virus injection, implantation of chronic window and holder

For in vivo 2-P calcium imaging, implantation of chronic window and holder was conducted while virus injection. 1.3 µL Ready-to-use pGP-AAV-syn-jGCaMP8f-WPRE AAV1 (Addgene, 162376, titer ≥ 7×10¹² vg/mL) was injected into primary visual cortex ((coordinates from Bregma: left 2.5 mm, AP -2.7 mm, Z 0.6-0.15 mm, 30°). Mice anesthetized with isoflurane in O2 (Abbvie, Ludwigshafen, Germany) were placed in a stereotactic frame (Kopf Instruments, CA, USA) and warmed with a heating pad (World Precision Instruments, Sarasota, FL, USA). For chronic window preparations, the skull was exposed by cutting off the scalp and well cleaned for fixing the head-holder. A round craniotomy with maximum diameter of 2.5 mm was carefully conducted over primary visual cortex (V1, left 2.5 mm, AP -2.7 mm). Diluted viral solution were injected via a custom-made tip-loading system, including a pulled micro-pipette (Hirschmann Laborgeräte, Eberstadt, Germany), a syringe and plastic tubing, using manual pressure. The pulled micro-pipette was slowly inserted into the exposed brain and approximately 300 nl of the viral solution was injected with an injection speed around 0.2 μl / min stepwise from 600 μm depth, targeting layer V / VI, to 200 μm, targeting layer II / III. Before retraction, the micro- pipette remained in place for 5 - 10 minutes helping the dispersion of virus solution in brain tissue meanwhile preventing an outward flow. After the virus injection, the opening was closed with a circular cover slip (Electron Microscopy Sciences, Hatfield, PA, USA) with a diameter of 3 mm and sealed onto the skull with surgical glue. The head holder was fixed onto the exposed skull with UV-glue (Polytec, Waldbronn, Germany). For a more detailed protocol please see ^38^.

### Two-photon imaging on awake mice

*In vivo* recordings were conducted using a custom-built 2-photon microscope equipped with a resonance scanner (LaVision Biotec, Bielefeld, Germany) and a Ti:Sapphire laser operating at 920 nm wavelength as light source (Coherent, Santa Clara, FL, USA). A 20x water immersion objective (Nikon) was used for imaging. Image acquisition was controlled by ImSpector Pro software (LaVision Biotec) at a frame rate of 30.9 Hz with a field of view of 458 x 458 μm^2^. Habituation of GCaMP8f injected mice with chronic window and holder were performed 7 days before the awake recording to adapt the awake head-fixing imaging system. Spontaneous 2- Photon calcium imaging was recorded for 15 min for each trial. Calcium transients were represented as relative changes in fluorescence (df/f).

### Visual stimulation by virtual reality

A drifting grating visual stimulation was presented in a virtual reality (VR) system consisting of a movement detectible air-float jet-ball system (PhenoSys, Berlin, Germany) and a 270° monitor system for delivering visual stimulus ^23^. The distance between the screens and eyes of the head-fixed mouse in the awake-behaving 2-P imaging system was approximately 50-60 cm. From this position, the screens covered the whole field of visual perception. Stimuli were conducted per cycles. In each stimulation sequence, 3-second grey screen and 2-second drifting gratings interleaved (Fig. 3A). Drifting gratings appeared randomly in 8 directions in each sequence to prevent the possibility of adaptation of mice to the stimulus (0°, 45°, 90°, 135°, 180°, 225°, 270°, 315°) ^39^. In each visual stimulation experiment, 20 stimulation sequences were performed.

### Two-photon Data Analysis

Data analysis was performed using a combination of CAIMAN ^40^ and a custom-built analysis pipeline. The process begins with the manual inspection of a few randomly selected recordings using a custom script to determine the optimal parameters for CAIMAN (e.g., patch size, overlap, number of components per patch). The same parameters are used for all recordings. Once optimized, raw recordings undergo motion correction, segmentation, and df/f trace extraction using CAIMAN. The resulting data is saved in an HDF5 file.

Next, a semi-automated pipeline loads these results into a custom application for automated calcium transient detection. Regions of interest (ROIs) with a signal-to-noise ratio (SNR) below 3 are excluded. To smooth the data, a 1D Gaussian filter is applied to the df/f traces. Transient peaks are then detected using scipy signal’s peak-finding function, applying a prominence and height threshold of at least 20%, with a minimum peak distance of 0.4 seconds (based on typical GCaMP8f transient in our datasets). This peak detection is run both on the filtered df/f traces and the denoised traces output by CAIMAN.

Transient onset and offset times are defined as the points where the signal drops to 90% of the peak height, within a time window guided by the calcium indicator’s half-time and peak height. Any transients lasting more than three times the expected duration, or with a decay time faster than their rise time, are discarded. ROIs with fewer than one transient per 15 minutes are labeled as inactive.

After this automated detection step, a manual quality control stage is used to exclude ROIs that show false positives (e.g., motion artifacts) or to re-include ROIs that were falsely excluded by the automated filters. This step does not involve reviewing individual transients but instead focuses on the overall quality and reliability of each ROI. The final set of detected transients, including their peak locations, heights, onsets, and offsets, is saved back to the same HDF5 file. To facilitate further analysis, we generate a binarized activity trace for each ROI. These traces are similar in format to df/f traces but are simplified to highlight periods of neural activity while minimizing noise. Binary traces retain the onset-to-peak portion of each detected transient while ignoring the decay phase, which is not informative about neuronal firing. Each transient segment is assigned a fixed value corresponding to its normalized df/f peak height, i.e. scaled between 0 and 1 across all detected events for that ROI. This results in a cleaner, binary representation of neural activity that serves as the basis for all downstream analyses. This binary activity matrix of all ROIs was then used to create pairwise correlations between each pair of ROI and was depicted in a heatmap as synchronicity.

### Synapse staining

Free-floating immunolabeling of Shank2 and GABAA was performed on perfusion fixed brains. Following transcranial perfusion with 4% PFA, brains were dissected and stored in 4% PFA overnight, followed by PBS with 0.01 sodium azide. Brains were cut in 40 µm coronal sections, using a vibratome, washed and subsequently unmasked in citric buffer pH6 (0.01 M citric acid in H20) at 85°C for 30 minutes. Sections were blocked in 10% normal goat serum (NGS) blocking solution (0.2% Triton X-100 in PBS), washed and then incubated with either 1:500 Shank2 (Thermo Fisher Scientific, Cat# PA5-78652) or 1:1000 GABAA (Synaptic Systems, Göttingen, Germany, Cat# 224003) primary antibody, both diluted in 3% NGS blocking solution overnight. The secondary antibody Alexa Fluor 647 goat anti-rabbit was diluted 1:1000 in 3% blocking solution und applied together with DAPI (1:1000) for one hour at room temperature. Sections were washed with PBS after incubation and subsequently mounted on slides. The slides were then dried at room temperature, covered with ProLong^TM^ Gold mounting media (Thermo Fisher Scientific) and capped with glass coverslips.

The synaptic stainings were captured using a confocal laser scanning microscope with a 63x immersion oil objective and a 4x digital zoom (1024 x 1024 pixels). The region of interest was determined as the visual cortex (left: 2.5 mm, AP -2.7 mm, Z 0.5 mm). For one pair (one animal, irradiated and control hemisphere) the microscope settings were kept constant and unchanged, in total six recordings were captured for each condition. Quantification of Shank2 and GABAA puncta was done by using the open-source image analysis software Image J. The background of one pair was determined and the mean value across all recordings was calculated. The threshold for one puncta was set to 2.5x the background intensity. The images were further processed by the despeckle and watershed function of image J, finally quantification was done by using the “Analyze particles” algorithm with a minimum particle size of 0.03 µm^2^.

### Statistics

Statistical analyses were performed using GraphPad Prism software. Kruskal-Wallis test was used. Data are presented with median and the box and whiskers plot. Box indicates the 25-75th percentile, whiskers indicate 10-90th percentile, and “+” indicates the mean.

## Acknowledgments

Acknowledgments follow the references and notes but are not numbered. Start with text that acknowledges non-author contributions and then complete each of the sections below as separate paragraphs.

## Author contributions

Conceptualization: AS, FR, JRA, MBS, MS

Methodology: TF, FL, FP, JM, SH, JA

Investigation: WF, FL, FP, JM, SH, JA

Visualization: TF, SA, FL, FP, JM

Funding acquisition: JRA, AS

Supervision: AS, MS

Writing – original draft: AS, WF, FR, JM, FL, FP, MS

Writing – review & editing: JRA, MS, HW, DC, SA, AS

## Data and materials availability

All data, and code is available upon request.

## Supplementary Figures

**Fig. S1.**
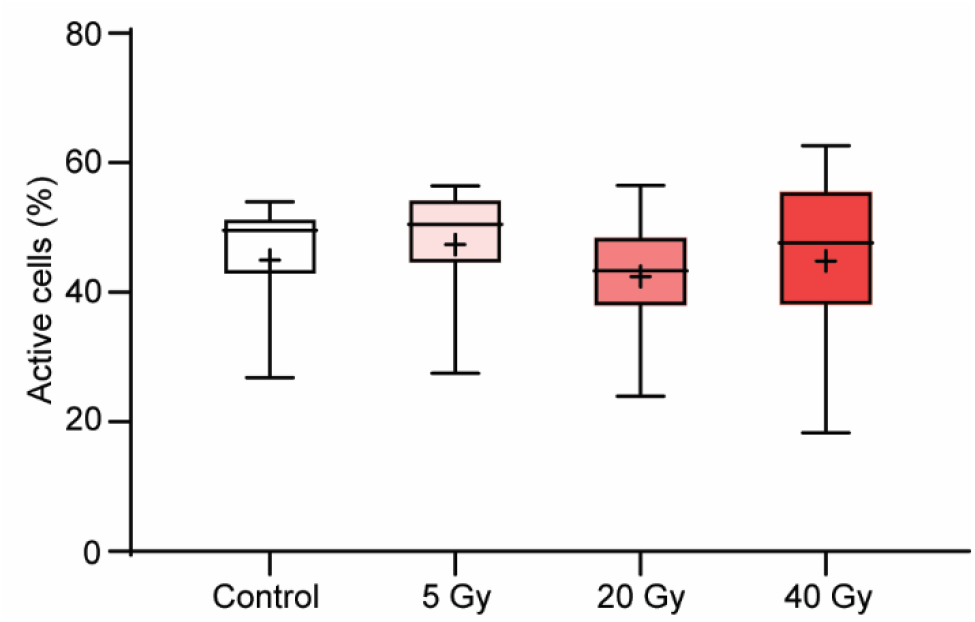
Irradiation did not affect the portion of active cells. Box and whiskers plot. Box indicates the 25-75th percentile, whiskers indicate 10-90th percentile, and “+” indicates the mean. Kruskal-Wallis test.

**Fig.S2.**
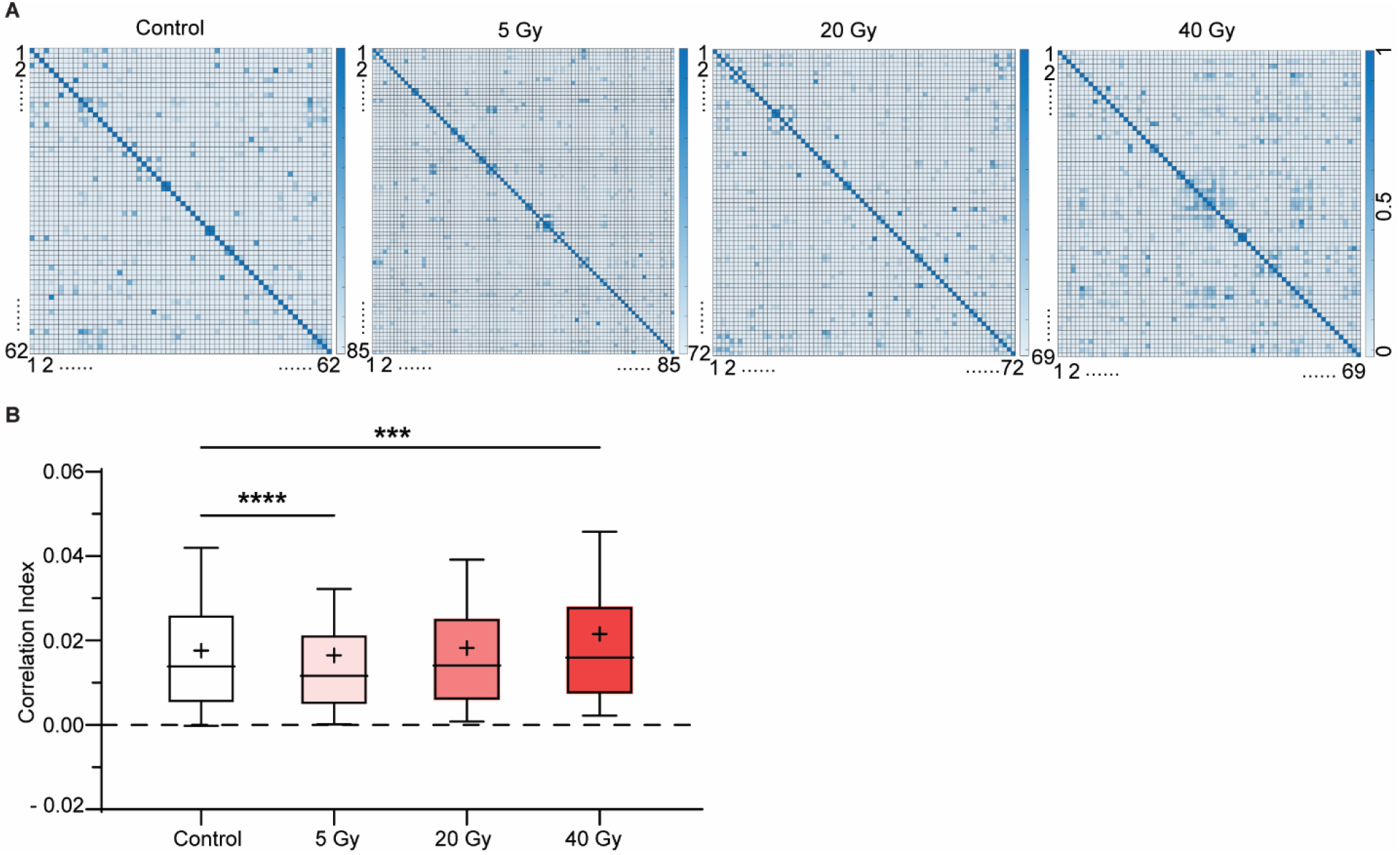
Impact of irradiation on network synchronization. (**A**) Representative heatmaps of the correlation index. (**B**) 5 Gy irradiation decreased the correlation index whereas 40 Gy irradiation increased the correlation index. Box and whiskers plot. Box indicates the 25- 75th percentile, whiskers indicate 10-90th percentile, and “+” indicates the mean. ***p < 0.001. ****p < 0.0001. Kruskal-Wallis test.

